# Neural activation down to the spinal cord during action language? A transcranial magnetic stimulation and peripheral nerve stimulation study

**DOI:** 10.1101/2025.04.02.646751

**Authors:** W. Dupont, N. Amiez, R. Palluel-Germain, A. Martin, M. Perrone-Bertolotti, C. Madden-Lombardi, F. Lebon

## Abstract

Language comprehension is increasingly recognized as extending beyond the traditional linguistic system to engage motor and perceptual processes. This perspective is supported by numerous studies demonstrating that understanding action-related words often induces behavioral and neurophysiological changes in the motor system. However, it remains unclear whether the influence of action language on the motor system is restricted to cortical regions, or whether it also extends to spinal structures, as observed during motor imagery. To address this, we used transcranial magnetic stimulation and peripheral nerve stimulation to assess corticospinal excitability and cortico-motoneuronal transmission, respectively. Fifteen healthy and right-handed volunteers participated in four conditions: (i) rest, (ii) kinesthetic motor imagery of finger and wrist flexion, (iii) reading action sentences, and (iv) reading non-action sentences. As anticipated, corticospinal excitability increased during both kinesthetic motor imagery and action reading compared to rest. Interestingly, while kinesthetic motor imagery also led to the expected increase in cortico-motoneuronal transmission, no such modulation occurred during action reading. These findings suggest that action reading do not modulate the excitability of high-threshold motoneurons at the spinal level, contrary to motor imagery. Further investigation is needed to test whether action reading activate lower-threshold spinal structures, such as interneurons involved in spinal pre-synaptic inhibition.

## 1. Introduction

For decades, researchers postulated that linguistic processing operated independently of perceptual and motor systems, implying a fundamental separation between cognition and bodily movements (Fodor, 1980; Pylyshyn, 1984). Nevertheless, language comprehension may extend beyond this traditional linguistic system to encompass motor and perceptual processes (Barsalou, 2008; Bechtold et al., 2023; Pulvermüller, 2005; Pulvermüller and Fadiga, 2010).

Along those lines, the comprehension of action words is often accompanied by changes in the motor system, in both behavioral and neurophysiological aspects (Andres et al., 2015; Gentilucci et al., 2000; Klepp et al., 2019; Pulvermüller et al., 2001; Rabahi et al., 2013, 2012; Taylor and Zwaan, 2008; Zwaan and Taylor, 2006). For example, neuroimaging studies demonstrate that understanding action-related words is associated with increased activation in motor areas of the brain, such as premotor and primary motor cortices (Aziz-Zadeh et al., 2006; Hauk et al., 2004; Tettamanti et al., 2005; Van Dam et al., 2010; Wu et al., 2013). Furthermore, this neural activation is closely tied to an increase in corticospinal excitability, as measured by motor-evoked potential (MEP) amplitude in Transcranial Magnetic Stimulation (TMS) studies (Dupont et al., 2025, 2022; Innocenti et al., 2014; Labruna et al., 2011; Papeo et al., 2009). Taken together, these studies indicate that understanding action words can directly modulate the functional state of the motor system (for reviews, see Madden-Lombardi and Dupont, 2023; Papeo et al., 2013).

Over the past several decades, a growing literature has associated these results with spontaneous and unconscious motor simulations, providing sensory-motor features of the described action, and consequently improving understanding of the action (Barsalou 1999; Glenberg and Kaschak 2002; Gallese and Lakoff 2005; Tomasino et al. 2008; Tomasino and Rumiati 2013; Dupont et al. 2024). These observations raise a fundamental question: could the influence of the comprehension of action words on our motor system extend beyond the cortical level? While this hypothesis may initially seem speculative, it aligns seamlessly with the well-established framework of motor imagery — a cognitive process defined as the explicit mental simulation of an action and its sensorimotor consequences, without concomitant movement. Like action word comprehension, motor imagery is associated with an activation of the motor network (Hardwick et al., 2018; Hétu et al., 2013; Lotze et al., 1999) and an increase in corticospinal excitability (Fadiga et al. 1998; Rossini et al. 1999; Tremblay et al. 2001; Facchini et al. 2002; Grosprêtre et al. 2016b; Dupont et al. 2024). Furthermore, recent research has highlighted the potential generation of a subliminal cortical output during motor imagery, suggesting that this cognitive process is not limited to the brain, but could eventually engage spinal structures. For instance, both motoneuronal excitability, reflected by the F-reflex, and cortico-motoneuronal transmission, measured by cervicomedullary-evoked potentials (CMEPs), increase during kinesthetic motor imagery (for F-reflex, see Bunno, 2019; Bunno et al., 2017; Fujisawa et al., 2011; Fukumoto et al., 2022; Hara et al., 2010; Suzuki et al., 2013; Taniguchi et al., 2008; for CMEPs, see Grosprêtre et al., 2016a).

Finally, growing evidence indicates that the neural mechanisms underlying action reading and motor imagery exhibit remarkable similarities, with the potential for reciprocal influence between these processes (Bonnet et al., 2022; Dupont et al., 2023; Gallese and Lakoff, 2005; Muraki et al., 2023; Yang and Shu, 2014). Both processes involve motor simulations associated not only with behavioral performance changes (e.g., reaction time, strength, or movement kinematics), but also with modulations of corticospinal excitability and motor cortex activity. In this regard, it remains unclear whether motor system activation generated by action reading is limited to cortical structures or if spinal activation is also engaged, as during motor imagery. Since MEP amplitude reflects the excitability of the entire corticospinal pathway, comparing changes in CMEP and MEP responses could help differentiate whether these neural modulations originate at the cortical or spinal level (Mcneil et al., 2013; Taylor et al., 2002). Therefore, the present research aims to shed new light on this issue by probing whether action reading, like motor imagery, involves the modulation of physiological measures at the spinal level. If action reading triggers a subliminal cortical output similar to motor imagery, we would observe an increase of both corticospinal excitability (MEP amplitude) and cortico-motoneuronal transmission (CMEP amplitude) in comparison to rest.

## 2. Material and methods

### 2.1. Participants

We recruited 15 right-handed individuals in this cross-sectional study (9 women, mean age: 23; range: 18-34). A study conducted with young healthy participants (Grosprêtre et al. 2016) demonstrated increased CMEP during motor imagery, compared to rest. Based on the effect size observed in that study (1.64), assuming a 5% alpha level and 80% power, 6 analyzable participants were required. However, considering that we expect a smaller effect size (0.82) for action reading compared to motor imagery, 15 participants were required. We ensured right handedness laterality with the Edinburgh inventory (Oldfield 1971; range: 0.64-1). All participants were French native speakers without neurological, psychological or physical disorders. They completed the questionnaire by Lefaucheur et al. (2011) to determine whether they were eligible for transcranial magnetic stimulation, and confirmed their participation by providing written consent. All procedures were in accordance with the Declaration of Helsinki and were approved by an ethics committee (excluding pre-registration; Comité de Protection des Personnes Nord-Ouest I, ClinicalTrials.gov Identifier: NCT06478303-01/08/2024).

### 2.2 Study design

Participants came to the laboratory for a single session. During the experiment, the participants were comfortably seated in a chair with their right hand in a neutral position, their right upper arm vertically aligned along the trunk, and the forearm semi-pronated and flexed at approximatively 90°. Using transcranial magnetic stimulation (TMS) and peripheral nerve stimulation (PNS), we elicited motor-evoked potentials (MEPs), cervicomedullary-evoked potentials (CMEPs) and maximal muscular responses (M_MAX_) in the right Flexor Carpi Radialis (FCR) muscle (see below for details). These measurements — recorded while participants remained still, i.e., without pre-muscular contractions — allowed us to investigate the corticospinal excitability, the cortico-motoneuronal transmission and the maximal muscle action potential at rest, during motor imagery, and during the silent reading of action and non-action sentences. The experiment was performed in a counterbalanced block design, with two blocks for each condition: Rest, Motor Imagery, Action Reading, and Non-Action Reading. Each condition included a TMS block with 16 MEPs, and an electrical stimulation block comprising 5 to 8 CMEPs (depending on the participant’s tolerance and the reproducibility of responses) and 2 M_MAX_. To control for potential order effects, the sequence of blocks was counterbalanced across participants. Volunteers read stimuli presented on a 19-inch LCD monitor via custom software, which synchronized TMS and PNS triggers and electromyographic recordings.

### 2.3 Tasks and stimuli

#### Action and non-action reading conditions

One hundred and four French sentences were generated; half referring to finger and wrist actions (e.g., “The pan is dirty, I scour it”; “La casserole est sale, je la récure”) and half describing non-actions (e.g., “The pan is dirty, I see it”; “La casserole est sale, je la vois”). All sentences in both of these action and non-action conditions were presented in the first-person present tense and were created so that the target verb occurred at the end of the sentence. This final pronoun-verb segment was presented alone on a subsequent screen. Each trial started with a fixation cross (300 ms), followed by a black screen (500 ms) and a sentential context (3000 ms), and then the pronoun and target verb (2000 ms, Figure 1). All stimuli order was counterbalanced.

**Figure 1.**
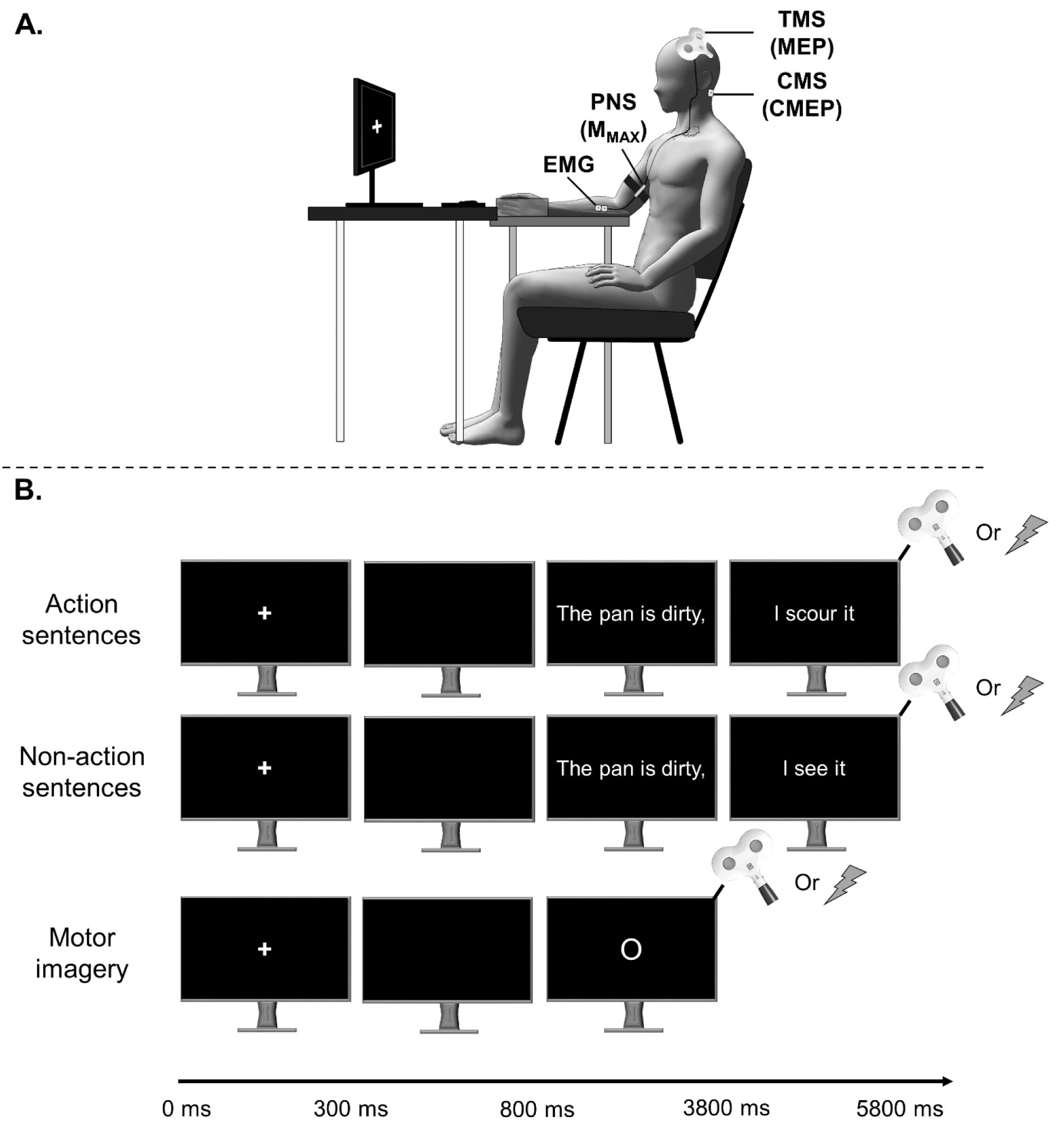
Experimental procedure. (**A**) Schematic illustration of the position of participants and stimulation devices. (**B**) Experimental procedure. Each trial started with an attentional fixation cross indicating the beginning of a trial. Action and non-action reading: the participants silently read sentences describing action or non-action concepts. We delivered TMS pulses and electrical stimulation at the optimal individual latency after the target verb presentation. Motor imagery: the participants imagined maximum voluntary contractions of wrist and finger flexion. We delivered TMS pulses and electrical stimulation 2000 ms after the appearance of the imagery onset (‘O’). TMS= transcranial magnetic stimulation; PNS= peripheral nerve stimulation; CMS= cervicomedullary stimulation; MEP= motor-evoked potentials; MMAX= maximal muscular response; CMEP= cervicomedullary-evoked potentials; EMG= electromyography.

Using the Lexique.org database (New et al., 2004), we controlled various psycholinguistic factors between action and non-actions verbs (e.g., written frequency, number of letters and syllables, spelling neighbors, number of homographs, and imageability). In addition, to confirm the distinction between action and non-action sentences, eight participants that did not participate to the TMS and CMS study were asked to rate on a scale from 1 to 5 whether the sentence described a manual action or not (1 = absolutely not a hand action, and 5 = entirely a hand action). Statistical verb analyses are detailed in Supplementary Table 1.

First, we conducted a series of practice TMS trials to determine each participant’s optimal latency for TMS, CMS and PNS administration during action and non-action reading (selecting from intervals of 200, 300, 400, or 500 ms after the target verb appeared). Since the delay between TMS/CMS stimulation and muscle response differs by only ∼2 ms across these modalities (Taylor, 2006), the same latency was applied for all. This latency was identified as the one eliciting the highest and most consistent MEP amplitude across 12 stimulations for both action and non-action reading conditions (Dupont et al., 2025). Once established, the participant’s individualized latency was used for the delivery of 16 TMS pulses (block TMS), along with 5 to 8 CMEP and 2 M_MAX_ (block electrical stimulations), following the presentation of the target verb in each condition.

#### Motor imagery condition

In the motor imagery condition, volunteers were asked to imagine maximum voluntary contractions of wrist and finger flexion movements held for 3 seconds in the kinesthetic modality. The instructions were as follows: “try to imagine yourself flexing your fingers and wrist, by feeling the sensations as if you were doing the movement”. TMS pulses were delivered 2000 ms after the appearance of the imagery signal (“O” on the screen). The inter-trial interval was 7000 ms. As in the reading task, 16 TMS pulses (block TMS), 5 to 8 CMEP and 2 M_MAX_ (block electrical stimulations) were administered during kinesthetic motor imagery.

#### Rest condition

Finally, participants received 16 TMS pulses (block TMS), along with 5 to 8 CMEP and 2 M_MAX_ (block electrical stimulations) at rest (during presentation of the fixation cross), which served as baseline.

### 2.4 Neurophysiological parameters

#### Electromyography recording

We recorded surface electromyography (EMG) via 10-mm-diameter surface electrodes (Contrôle Graphique Médical, Brice Comte-Robert, France) positioned over the right Flexor Carpi Radialis (FCR) muscle. Before positioning the electrodes, the skin was shaved and cleaned to reduce EMG signal noise. The EMG signal was amplified with a bandpass filter (10-500 Hz) and then sampled at 5000 Hz (AcqKnowledge; Biopac Systems, Inc., Goleta, CA). We computed the root mean square EMG signal (EMGrms) 100 ms prior to the stimulus artefact to ensure that MEP, CMEP and M_MAX_ were not biased by muscle pre-activation during action reading and motor imagery in comparison to rest (see Supplementary Table 3).

#### Transcranial magnetic stimulation

Using a figure-eight coil (70 mm diameter) connected to a Magstim 200 stimulator (Magstim Company Ltd, Whitland), we administered single-pulse TMS over the motor area of the right FCR muscle. The coil rested tangentially to the scalp with the handle directed backward and laterally at a 45° angle from the midline. Using a neuronavigation system (BrainSight, Rogue Research Inc.), the muscle hotspot was identified as the location eliciting the highest and most consistent MEP amplitude for the FCR muscle at rest. This hotspot was pinpointed through a regular grid of 4 by 4 coil positions with a spacing of 0.5 cm (centered above the FCR cortical area x=-44.7, y=-13.8, z=85.7; Négyesi et al. 2020; Sondergaard et al. 2021). Then, the resting motor threshold was estimated for each individual, corresponding to the minimal TMS intensity required to evoke MEPs of 50µV peak-to-peak amplitude in the right FCR muscle for 5 out of 10 trials (Rossini et al., 2015). The intensity of TMS pulses was set at 120% of the resting motor threshold (mean intensity: 46 ±10.39 % of maximal stimulator output, range: 34-67%; Mean MEP amplitude at rest: 0.29 ±0.22 mV).

#### Electrical stimulation

##### Cervico-medullary stimulation

In order to probe whether a subliminal motor output extended beyond the brain and reaches spinal structures, we recorded CMEP in the FCR muscle. This approach employed cervicomedullary stimulation to directly probe the cortico-motoneuronal transmission by generating a single volley in descending axons at the pyramidal decussation level (Mcneil et al., 2013; Taylor, 2006). Interestingly, CMEP primarily exhibits a monosynaptic component without influence of presynaptic inhibition. Cervicomedullary stimulations were administered using a DS7R constant current research stimulator (Digitimer, Hertfordshire, UK) with 200 µs-width pulses, via two AgCl electrodes positioned 1-2cm posterior to the superior tips of the mastoid processes (left cathode and right anode). The cervico-medullary stimulation intensity was set to match the MEP response at rest with the TMS intensity set at 120% of the resting motor threshold (mean intensity: 123 ±21.19 mA, range 99-170 mA; mean CMEP amplitude at rest: 0.23 ±0.12 mV). Interestingly, alpha motoneurons targeted by this technique largely overlap with those of MEP (Mcneil et al., 2013; Taylor, 2006; Taylor et al., 2002), as long as the MEP and CMEP amplitude is matched. We ensured that MEP and CMEP amplitude at rest recruited the same amount of motor units for each participant, characterized by the percentage of M_MAX_ amplitude (see Supplementary Table 2).

##### Median nerve stimulation

To evoke M_MAX_ in the FCR muscle, we employed the same constant current stimulator (Digitimer DS7R) to deliver 1 ms-width monophasic rectangular pulses over the median nerve. Electrodes for stimulation were meticulously positioned along the median nerve, close to the cubital fossa, just below the muscle belly of the biceps, with the anode distally positioned. After securing the stimulation electrodes with straps, we determined the stimulation intensity eliciting the maximum response amplitude (M_MAX_) using a recruitment curve initially incremented by 2 mA, and then adjusted to 1 mA when nearing the maximum response. To ensure supramaximal stimulation, the intensity was increased to 130% (mean intensity: 14.24 ±5.45 mA, range 6-27 mA; mean M_MAX_ amplitude: 10.8 ±4.63 mA). Subsequently, the M_MAX_ was measured in each experimental condition and used to normalize the MEPs (MEP/M_MAX_) and CMEPs (CMEP/M_MAX_).

### 2.5 Data and statistical analysis

#### Data extraction and analysis

Matlab (The MathWorks, Natick, Massachusetts, USA) was employed to extract EMG and we measured peak-to-peak M_MAX_, CMEP and MEP amplitude. Then, we discarded outliers falling outside the range of +/-2 SDs from individual means for each condition (5.57% of MEP data and 5.29% of CMEP data). MEP and CMEP peak-to-peak amplitude was normalized to the mean M_MAX_ (%M_MAX_) obtained within each block, ensuring control over potential changes in peripheral neuromuscular excitability and allowing interindividual comparison.

#### Statistical analysis

Normality and sphericity of the data were evaluated with Shapiro-Wilk and Mauchly tests, respectively. Given that the assumptions of normality and sphericity were violated, non-parametric statistical tests were employed throughout the analysis.

To analyze modulations of corticospinal excitability (MEP/M_MAX_), cortico-motoneuronal transmission (CMEP/M_MAX_), and M_MAX_, separate Friedman ANOVAs were conducted for each dependent variable, with Condition (rest, motor imagery, action reading, non-action reading) as a within-subject factor. If the Friedman ANOVAs yielded significant results, post hoc analyses were performed using multiple paired Wilcoxon signed-rank tests to compare across all conditions (for each dependent variable). Finally, Bayesian factor analyses with a prior Cauchly scale of .707 (Morey and Rouder, 2011) were performed to complete our interpretation, offering a measure of the strength of evidence in support of either the alternative or the null hypothesis. To illustrate, a BF₁₀ of 1 indicates no evidence to support either hypothesis, whereas a BF₁₀ between 1 and 3, or between 3 and 10, provides marginal and moderate support for the alternative hypothesis, respectively.

We also applied Friedman ANOVAs to compare the EMGrms before the stimulation artefact in all conditions (see Supplementary Table 3) to ensure that MEP, CMEP and M_MAX_ amplitudes were not biased by muscle activation preceding stimulation. Furthermore, we applied Friedman ANOVAs to compare the target positioning, angular alignment, and twist of the TMS coil across conditions, to ensure that MEP amplitude was not biased by variations in TMS coil placement relative to the hotspot (see Supplementary Tables 4 and 5).

We performed statistics with Statistica (Stat Soft, France) and JASP software, and the data are presented with mean values (±SD). The alpha value was set at .05. For paired sample Wilcoxon tests, Cohen’s d thresholds for small, moderate, and large effects were set at .2, .5, and .8, respectively (Cohen, 1988). All post hoc analyses were performed using Benjamini and Hochberg correction (Benjamini and Hochberg, 1995), which adjusts significance thresholds according to the number of comparisons, and the ranked p-values in ascending order: .01, .02, .03, .04, and .05.

## 3. Results

### 3.1. M_MAX_ amplitude

No main effect of Condition was found [Friedman ANOVA: r(3) = –.016, p = .510], suggesting no significant differences in M_MAX_ amplitude among rest (11.07 ±5.03 mV), kinesthetic motor imagery (11.04 ±5.05 mV), action (11.10 ±5.09 mV) and non-action reading (11.08 ±5.05 mV). While MEP and CMEP were normalized to M_MAX_ (%M_MAX_) to control for potential changes in peripheral neuromuscular excitability demonstrating no change in muscular excitability across conditions, this enable to probe the proportion of motor units activated by TMS and CMS.

### 3.2. MEP amplitude – Corticospinal excitability

We observed a main effect of Condition [Friedman ANOVA: *r*(3) = 0.35, *p* < .001] with greater MEP amplitude for kinesthetic motor imagery (5.08 ±3.35 %M_MAX_; see Table 1)) and action reading (4.08 ±3.02 %M_MAX_) compared to rest (3.41 ±2.76 %M_MAX_). Additionally, MEP amplitude during action reading was significantly higher than during non-action reading (3.14 ±2.59 %M_MAX_), which did not differ from rest. Finally, no significant difference in MEP amplitude was found between kinesthetic motor imagery and action reading (Fig.2). Bayesian factor analysis provides moderate evidence supporting the alternative hypothesis for both kinesthetic motor imagery and action reading, strengthening the idea of significantly increased MEP amplitude in these conditions compared to rest.

**Figure 2.**
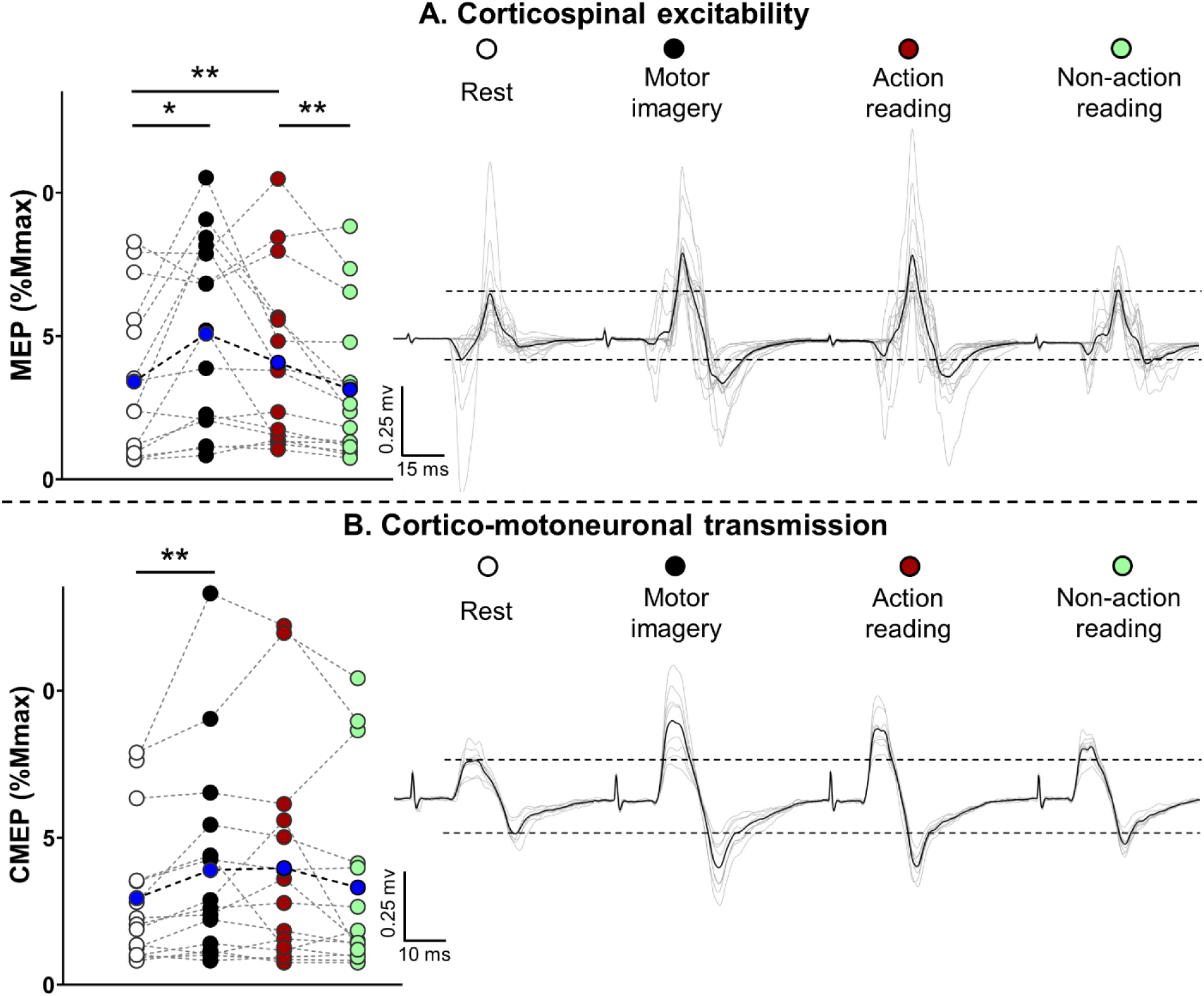
MEP (**A**) and CMEP (**B**) amplitude expressed as a percentage of maximal muscular response (%M_MAX_) at rest, during motor imagery, and during the silent reading of action and non-action sentences. The left side of the figure displays individual data points, with each condition represented by a unique color. Blue circles indicate the mean value for each condition. The right side of the panel illustrates the MEP and CMEP modulations. The asterisks denote statistically significant p-values after applying the Benjamini-Hochberg correction.

**Table 1.** Statistical values for corticospinal excitability.

| Comparisons | *P*<br>values | Corrected<br>threshold | Bayes<br>factor<sub>10</sub> | Cohen's<br>*d* |
| --- | --- | --- | --- | --- |
| Action reading vs. Non-action reading | .002* | .01 | 7.859 | .823 |
| Action reading vs. Rest | .008* | .02 | 3.179 | .678 |
| Motor imagery vs. Rest | .020* | .03 | 3.735 | .705 |
| Action reading vs. Motor imagery | .280 | .04 | .754 | .416 |
| Non-action reading vs. Rest | .460 | .05 | .492 | .315 |

*Note: An asterisk (\*) indicates P-values that are significant according to the corrected threshold with the Benjamini-Hochberg procedure.*

### 3.3. CMEP amplitude – Cortico-motoneuronal transmission

We observed a main effect of Condition [Friedman ANOVA: *r*(3) = 0.116, *p* = .048] with greater CMEP amplitude during kinesthetic motor imagery (3.90 ±3.51 %M_MAX_) compared to rest (2.95 ±2.43 %M_MAX_; see Table 2). However, CMEP amplitude did not differ significantly between rest, action reading (3.97 ±3.75 %M_MAX_), and non-action reading (3.31 ±3.31 %M_MAX_, Fig.2). Furthermore, Bayesian factor analysis provides marginal evidence supporting the alternative hypothesis for both kinesthetic motor imagery and action reading, suggesting that CMEP amplitude might be increased during motor imagery, and to a lesser extent, during action reading.

**Table 2.** Statistical values for cortico-motoneuronal transmission.

| Comparisons | *P*<br>values | Corrected<br>threshold | Bayes<br>factor<sub>10</sub> | Cohen's<br>*d* |
| --- | --- | --- | --- | --- |
| Motor imagery vs. Rest | .004* | .01 | 2.435 | .634 |
| Action reading vs. Rest | .036 | .02 | 1.487 | .548 |
| Action reading vs. non-action reading | .088 | .03 | .708 | .402 |
| Non-action reading vs. Rest | .211 | .04 | .438 | .283 |
| Motor imagery vs. Action reading | .691 | .05 | .267 | .052 |

*Note: An asterisk (\*) indicates P-values that are significant according to the corrected threshold with the Benjamini-Hochberg procedure.*

## 4. Discussion

The present study presents several noteworthy findings regarding motor simulations, whether elicited by motor imagery or action reading. First, we successfully replicated the well-established observation that motor imagery leads to a marked increase in corticospinal excitability compared to rest. Similarly, action-related language also resulted in greater corticospinal excitability compared to rest and to non-action reading (which did not differ from rest). Second, we found that motoneuronal excitability was significantly enhanced only during kinesthetic motor imagery relative to rest.

Regarding the first point, we confirmed the increase in corticospinal excitability, by means of MEP amplitude changes during kinesthetic motor imagery compared to rest (Dupont et al., 2024b, 2023; Facchini et al., 2002; Fadiga et al., 1998; Grosprêtre et al., 2016b; Kasai et al., 1997; Lebon et al., 2012; Rossini et al., 1999; Tremblay et al., 2001). In a similar vein, we observed that action reading — when compared to rest and non-action sentences — also resulted in an enhancement of corticospinal excitability (for similar results, see Dupont et al., 2025, 2024a, 2022; Innocenti et al., 2014; Labruna et al., 2011; for reviews, see Madden-Lombardi and Dupont, 2023). Thus, motor simulations generated during motor imagery and action reading may influence the excitability of the corticospinal pathway.

Regarding the second point, our findings provide further support for previous research demonstrating modulations at the motoneuronal level during kinesthetic motor imagery. More specifically, our results align with a previous study reporting that cortico-motoneuronal transmission increases during kinesthetic motor imagery, as measured by CMEP amplitude (Grosprêtre et al., 2016a). These findings lend further credence to the idea of a potential subliminal cortical output, suggesting that motor imagery may extend beyond the brain, potentially engaging subcortical and even spinal structures (Grosprêtre et al. 2016a; Grosprêtre et al. 2019). While the striking similarity between motor imagery and action reading in terms of corticospinal excitability is noteworthy, it is important to highlight that action reading did not significantly increase motoneuronal excitability. At first glance, this observation might indicate that the increase in corticospinal excitability triggered during action reading is likely primarily cortical in origin, rather than being driven by subcortical or spinal structures.

This absence of motoneuronal changes during action reading can be interpreted from two perspectives. First, the simulation of concepts evoked by action language may be restricted to the brain, without engaging the subliminal cortical output that reaches spinal structures, potentially indicating distinct underlying mechanisms compared to motor imagery. Second, one could argue that a subliminal motor output may still be present but to a lesser extent — more subliminal than during motor imagery — making it more challenging to activate high-threshold motoneurons. In other words, the motor simulation can be evoked at varying levels of richness and intensity, with motor imagery at a higher end of this continuum than action language. Indeed, although we did not observe a significant difference in MEP amplitude between motor imagery and action reading, previous studies have shown that MEP amplitude during motor imagery is greater when imagining a 60% force level compared to a 10% force level (Helm et al., 2015; Mizuguchi et al., 2013). While participants were instructed to imagine maximal contractions (100% force level) during motor imagery, the actions described in the sentences did not necessarily imply the same intensity. This difference in force intensity might explain the statistically non-significant increase in CMEP during action reading. To explore this hypothesis further, it would be interesting to investigate spinal structures with lower activation thresholds than motoneurons, such as presynaptic interneurons, which could be more responsive to this subtler activation triggered by action reading. For instance, recent studies have highlighted that imagined foot movement modulated the activity of interneurons involved in spinal presynaptic inhibition (Grosprêtre et al. 2016a; Grosprêtre et al. 2019). These interneurons — with activation thresholds lower than those of motoneurons — may be more susceptible to the subliminal activation potentially induced by action reading.

This study has some limitations that should be considered. First, the relative peak-to-peak amplitude, expressed as a percentage of the M_MAX_ wave for each stimulation, differed across participants, suggesting that the proportion of alpha motoneurons recruited was not consistent among individuals. However, the fact that we matched MEP and CMEP amplitude at rest, thereby recruiting the same amount of motor units for each participant during TMS blocks (range MEP amplitude at rest: 0.10-0.77 mV; mean MEP amplitude at rest: 0.29 mV) and CMS blocks (range CMEP amplitude at rest: 0.11-0.52 mV; mean CMEP amplitude at rest: 0.23 mV), provides reassurance regarding this variability. Lastly, a limitation concerning the linguistic stimuli arises from the fact that some sentences describing hand actions focused more on finger flexion rather than wrist flexion, which is the optimal movement for engaging the FCR muscle. Consequently, we speculate that these sentences may have resulted in less pronounced increases in MEP and CMEP amplitude following action reading.

To conclude, our results shed significant insights into the effects of motor imagery and action reading on both cortical and spinal structures. They support the notion that motor simulations involved in motor imagery may trigger a subliminal motor output initiated at the cortical level, with the potential to influence spinal activities. Furthermore, our results suggest that the impact of action words comprehension on the motor system may be restricted to cortical aspects, without engaging the subliminal cortical output that reaches spinal structures. Nonetheless, further investigations are needed to explore spinal structures with lower activation thresholds than motoneurons, as these may be more responsive to a subtler subliminal activation triggered by action reading.

## Data availability

All data and linguistic items from this study are available at https://osf.io/t75fr/?view_only=a47a7f9125df4e0f8c33ed86acfcdf8e

## Author contributions

L.F., M-L.C., P-B.M. and P-G.R. funding acquisition; D.W. and A.N. designed research; D.W. and A.N. recorded data; D.W. analyzed data; D.W. wrote the draft of the paper; L.F., M-L.C., M.A, P-G.R. and P-B.M. supervised research; A.N., P-G.R., A.M., P-B.M., M-L.C. and L.F. edited the paper.

## Acknowledgements

The authors are very grateful to Cyril Sirandré for help with the custom Neurostim software and the experimental setup.

## Supplementary section

**Table S1:**
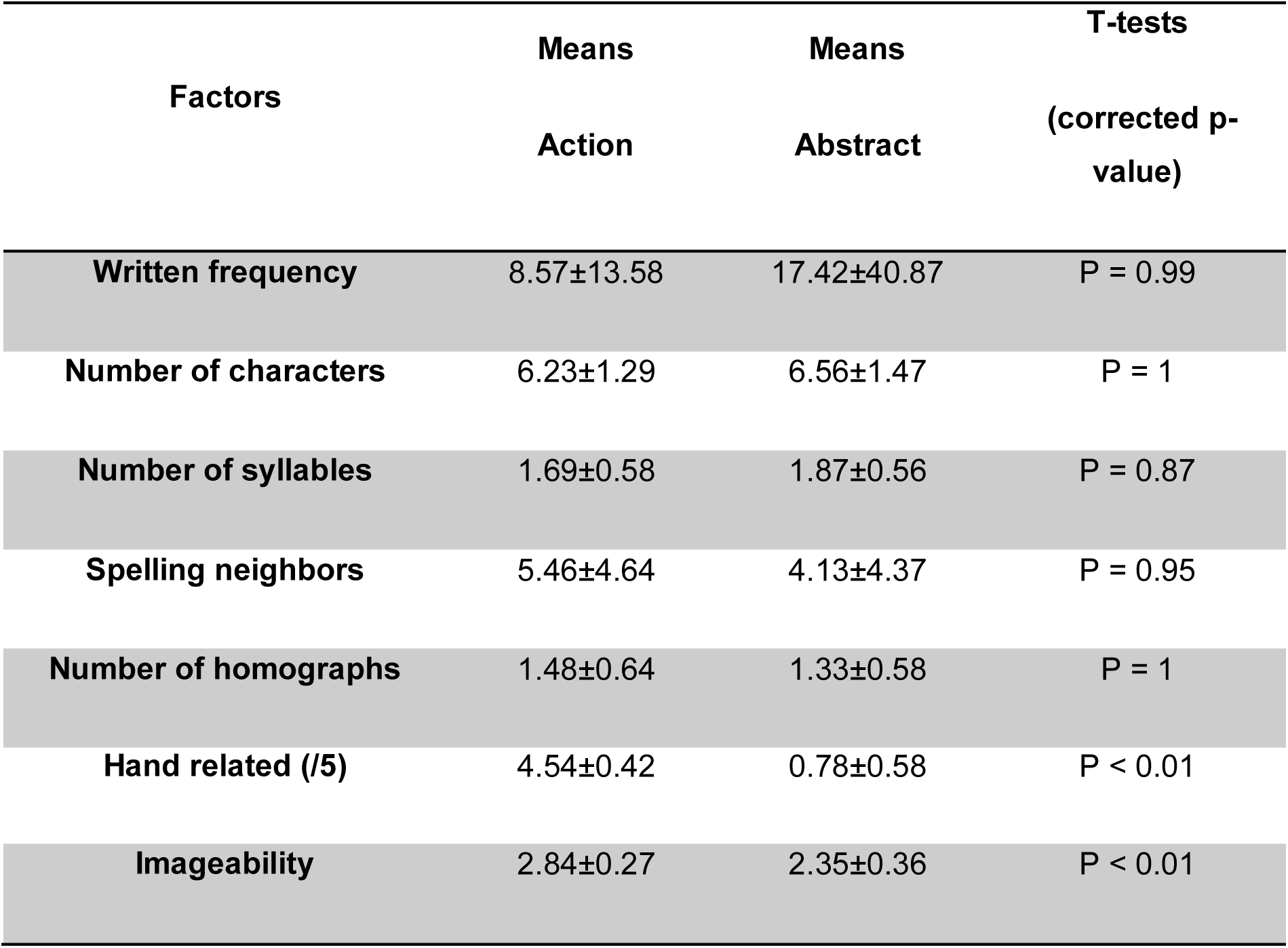
T-test results concerning the linguistic and psycholinguistic characteristics of stimuli.

| Factors | Means | Means | T-tests |
| --- | --- | --- | --- |
|  | Action | Abstract | (corrected p-value) |
| Written frequency | 8.57±13.58 | 17.42±40.87 | P = 0.99 |
| Number of characters | 6.23±1.29 | 6.56±1.47 | P = 1 |
| Number of syllables | 1.69±0.58 | 1.87±0.56 | P = 0.87 |
| Spelling neighbors | 5.46±4.64 | 4.13±4.37 | P = 0.95 |
| Number of homographs | 1.48±0.64 | 1.33±0.58 | P = 1 |
| Hand related (/5) | 4.54±0.42 | 0.78±0.58 | P < 0.01 |
| Imageability | 2.84±0.27 | 2.35±0.36 | P < 0.01 |

**Table S2:**
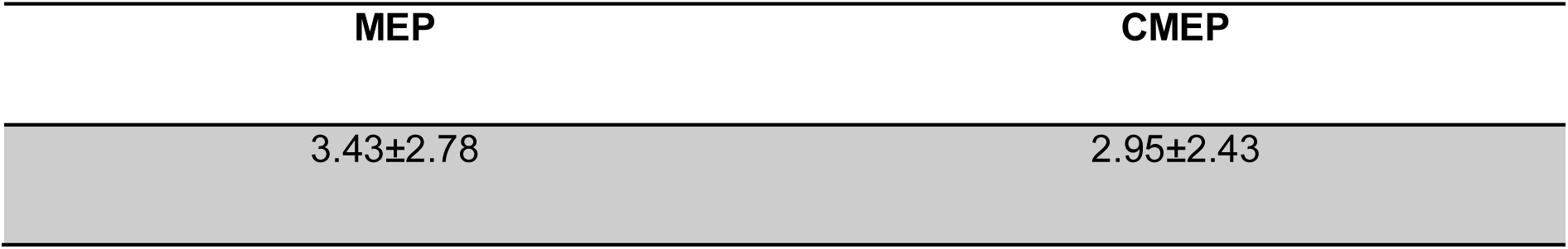
Comparison of the MEP and CMEP (%M_MAX_) at rest. A paired Wilcoxon signed-rank confirmed that the MEP and CMEP recruited the same amount of motor units (*t* = .722, *p* = .481).

**Table S3:**
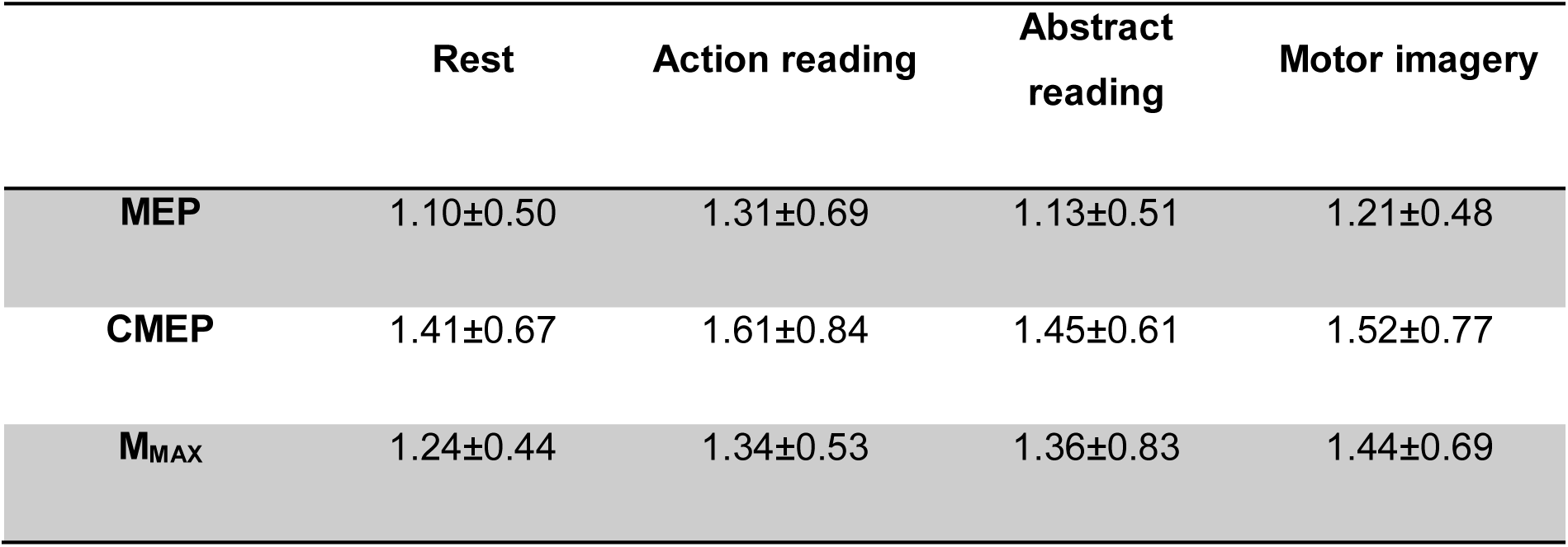
EMGrms activity (mean ±SD) in µV recorded for the Flexor Carpi Radialis before the TMS, CMS and M_MAX_ artifact for each condition (window of 200ms prior the artifact). The absence of muscular pre-activity was confirmed by separate Friedman ANOVAs revealing no significant difference in EMGrms before the MEP (*p* = 0.118; *r* = 0.069), CMEP (*p* = 0.313; *r* = 0.013), or M_MAX_ (*p* = 0.114; *r* = 0.070) between rest, motor imagery, action and non-action reading conditions.

|  | Rest | Action reading | Abstract reading | Motor imagery |
| --- | --- | --- | --- | --- |
| **MEP** | 1.10±0.50 | 1.31±0.69 | 1.13±0.51 | 1.21±0.48 |
| **CMEP** | 1.41±0.67 | 1.61±0.84 | 1.45±0.61 | 1.52±0.77 |
| **M<sub>MAX</sub>** | 1.24±0.44 | 1.34±0.53 | 1.36±0.83 | 1.44±0.69 |

**Table S4:** Differences in target positioning (mm), angular alignment (°), and twist (°) of the TMS coil relative to the hotspot were assessed for each condition. Separate Friedman ANOVAs confirmed the absence of significant coil movement, revealing no significant differences in target positioning (*p* = 0.379; *r* = 0.001), angular alignment (*p* = 0.240; *r* = 0.029), or twist (*p* = 0.379; *r* = 0.002) of the TMS coil across conditions.

|  | Rest | Action reading | Abstract reading | Motor imagery |
| --- | --- | --- | --- | --- |
| **Target positioning** | 0.35±0.06 | 0.34±0.07 | 0.33±0.07 | 0.32±0.06 |
| **Angular alignment** | 0.68±0.19 | 0.58±0.23 | 0.60±0.18 | 0.57±0.23 |
| **Twist** | -0.16±0.68 | -0.04±0.61 | -0.24±0.68 | -0.18±0.51 |

**Table S5:** MNI coordinates (x, y, z) of TMS stimulations localization averaged for each participant.

|  | X | Y | Z |
| --- | --- | --- | --- |
| **S01** | -36.830 | -8.721 | 67.085 |
| **S02** | -33.727 | -11.493 | 69.690 |
| **S03** | -33.705 | -11.651 | 69.745 |
| **S04** | -39.969 | -12.696 | 66.306 |
| **S05** | -39.949 | -12.769 | 66.346 |
| **S06** | -39.846 | -12.678 | 66.388 |
| **S07** | -36.914 | -15.605 | 68.902 |
| **S08** | -36.965 | -8.913 | 67.060 |
| **S09** | -37.110 | -8.697 | 67.284 |
| **S10** | -40.006 | -12.765 | 66.305 |
| **S11** | -40.095 | -12.736 | 66.231 |
| **S12** | -33.913 | -11.772 | 69.662 |
| **S13** | -36.995 | -15.875 | 68.901 |
| **S14** | -37.108 | -8.933 | 66.974 |
| **S15** | -37.028 | -15.808 | 68.868 |

